# Microbial community-based single-cell protein supports partial fishmeal replacement in juvenile Asian seabass diets across feeding trials and production scales

**DOI:** 10.64898/2026.06.17.732826

**Authors:** Ezequiel Santillan, Poh Leong Loo, Fanny Yasumaru, Hui Xu, Soheil Asgari Neshat, Ramanujam Srinivasan Vethathirri, Yan Zhou, Diana Chan Pek Sian, Stefan Wuertz

**Affiliations:** Singapore Centre for Environmental Life Sciences Engineering, Nanyang Technological University, Singapore 637551, Singapore; Aquaculture Innovation Centre, Temasek Polytechnic, Singapore 529757, Singapore; Advanced Environmental Biotechnology Centre (AEBC), Nanyang Environment & Water Research Institute (NEWRI), Nanyang Technological University, 1 Cleantech Loop, Clean Tech One, Singapore 637141, Singapore; School of Civil and Environmental Engineering, Nanyang Technological University, Singapore 639798, Singapore

**Keywords:** Asian seabass, single-cell protein, alternative aquafeeds, fishmeal replacement, microbial protein, sustainable aquaculture feed, circular bioeconomy

## Abstract

The growing demand for sustainable aquafeeds has intensified interest in alternative protein ingredients capable of reducing reliance on fishmeal without compromising fish performance. Here, we evaluated microbial community-based single-cell protein (SCP) as a fishmeal substitute in juvenile Asian seabass (*Lates calcarifer*) diets in two independent feeding trials of juvenile fish conducted over 49 and 56 days, respectively and compared them to a previous study that lasted 24 days. SCP was produced from nutrient-rich soybean-processing side streams by microbial communities in fermenters and incorporated into experimental diets at inclusion levels ranging from 10% to 100% fishmeal replacement. In the 24-day trial, a diet containing 50% fishmeal replacement with lab-scale produced SCP achieved 100% survival and a feed conversion ratio (FCR), specific growth rate (SGR), and weight gain comparable to the fishmeal control diet. In the 49-day trial using pilot-scale produced SCP, a 50% fishmeal replacement also maintained an FCR and feed intake comparable to the control, whereas complete replacement reduced feed intake and growth performance. In a 56-day pilot-scale trial that used 500-L fish tanks, diets containing up to 50% fishmeal replacement maintained comparable survival, weight gain, and SGR, although moderately higher FCR values were observed at higher SCP inclusion levels. Proximate composition and essential amino acid profiles of fish fed control or SCP-containing diets were comparable. Genome-resolved metagenomic analyses revealed diverse microbial taxa associated with the SCP. Collectively, these findings support microbial community-based SCP as a scalable and reproducible alternative protein platform for aquaculture feeds across independent trials and production scales.

## Introduction

Aquaculture is one of the fastest-growing food production sectors globally and plays an increasingly important role in food security and the supply of high-quality animal protein. However, continued aquaculture expansion depends heavily on the availability of sustainable and nutritionally adequate feed ingredients (Jones et al., 2020). Fishmeal remains one of the most widely used protein sources in aquafeeds due to its favorable amino acid composition, palatability, digestibility, and nutrient density (FAO-UN, 2020). Nevertheless, concerns regarding fluctuating prices, finite marine resources, and pressure on wild fish stocks have intensified efforts to identify sustainable alternative protein ingredients for aquaculture feeds (Glencross et al., 2024; Tacon, Metian, and McNevin, 2022). Recent analyses further emphasize the need to diversify aquafeed ingredient portfolios using scalable, lower-footprint protein sources capable of supporting continued industry growth (HatchBlue, 2024).

Single-cell protein (SCP), derived from microbial biomass, has emerged as a promising alternative protein source for aquaculture nutrition. SCP production systems offer several advantages, including rapid biomass generation, low land requirements, and the ability to utilize nutrient-rich side streams and agro-industrial side streams as substrates (Choi, Jung, and Lee, 2024). Such systems may simultaneously support nutrient recovery, waste valorization, and circular bioeconomy strategies for food and feed production (D’Almeida & de Albuquerque, 2025; Gao et al., 2025). Current SCP platforms include methanotrophic bacteria, purple non-sulfur bacteria, yeasts, filamentous fungi, hydrogen-oxidizing bacteria, and microalgae cultivated on substrates ranging from methane and methanol to food-processing by-products and side streams (Gao et al., 2025; Li et al., 2024). However, microbial groups and production systems differ substantially in nutritional composition, digestibility, functional properties, and production characteristics, resulting in variable aquaculture outcomes depending on microbial source, cultivation substrate, downstream processing, dietary formulation, and target species (D’Almeida & de Albuquerque, 2025; Mugwanya, Kimera, and Sewilam, 2026).

Asian seabass (*Lates calcarifer*) is a major aquaculture species cultivated throughout the Asia-Pacific region and represents an important candidate for evaluating sustainable alternative feed ingredients. Previous studies demonstrated that microbial proteins can partially replace fishmeal in Asian seabass diets, although outcomes vary substantially depending on SCP type and formulation. Enriched purple phototrophic bacterial biomass supported partial fishmeal replacement, whereas complete replacement negatively affected growth performance and feed conversion ratios (Delamare-Deboutteville et al., 2019). In contrast, methanotrophic bacterial SCP replacing 25–50% of fishmeal in barramundi fry diets resulted in reduced growth, lower survival, enteropathy, liver pathology, and gut microbiome dysbiosis (Samsing et al., 2024). Studies in other aquaculture species, including Pacific white shrimp (*Litopenaeus vannamei*), further support the potential of microbial proteins as alternative feed ingredients (Alloul et al., 2021; Felix et al., 2023; Santillan et al., 2025). Collectively, these findings indicate that SCP performance depends strongly on production system, microbial composition, and dietary formulation (HatchBlue, 2024; Mugwanya, Kimera, and Sewilam, 2026).

Most aquaculture SCP studies have focused on axenic or defined microbial production systems. In contrast, SCP generated through non-axenic microbial communities remains comparatively underexplored despite potential advantages in substrate flexibility, process robustness, and scalability (Vethathirri, Santillan, and Wuertz, 2021). Recent work demonstrated the feasibility of producing microbial community-based SCP from soybean-processing side streams using fermenters enriched in aerobic heterotrophic bacteria (Vethathirri et al., 2023; Vethathirri et al., 2025). The resulting biomass exhibited nutritionally relevant amino acid profiles, and a previous short-term feeding trial demonstrated successful 50% fishmeal replacement in juvenile Asian seabass diets without compromising growth performance or survival (Santillan et al., 2024). The present study further evaluated microbial community-based SCP diets across additional feeding trials conducted under longer growth periods, broader dietary inclusion levels, and progressively scaled aquaculture conditions. Fish growth performance, feed utilization, survival, and nutritional composition were evaluated across trials, and genome-resolved metagenomic analyses were performed to characterize the microbial communities associated with the SCP ingredient.

## Materials and methods

### Fish husbandry and culture conditions

Juvenile Asian seabass (*Lates calcarifer*; barramundi) were obtained from local commercial hatcheries (Allegro Aqua for Trial 1; Barramundi Group for Trials 2 and 3) and acclimated to laboratory conditions prior to the experiments. Fish were initially fed a commercial seabass diet (43% protein; Uni-President, Vietnam) during the acclimation period. Experimental fish were maintained under aerated brackish water conditions in recirculating aquaculture systems (RAS) or pilot-scale tank systems depending on the trial. Water quality parameters, including temperature, dissolved oxygen (DO), salinity, pH, ammonia, nitrite, and nitrate, were monitored throughout the study and maintained within acceptable ranges for juvenile Asian seabass culture.

In an earlier study (henceforth referred to as Trial 1), a 24-day feeding trial (Santillan et al., 2024) was conducted with SCP that had been produced under laboratory-scale conditions (fermenter size of 4 L). Briefly, juvenile seabass (19.7 ± 2.6 g) were randomly distributed into rectangular glass aquaria ( 120-L water volume) at a stocking density of 20 fish per tank, with three replicate tanks assigned to each dietary treatment (n = 3). The aquaria were connected to RAS. Water temperature ranged from 27 to 29 °C, DO remained above 4 mg L ^1^, salinity was maintained at 15 ppt, pH ranged from 7.0 to 7.9, and ammonia and nitrite concentrations remained below 0.5 ppm.

In the present study, two independent feeding trials were performed over longer growth durations than that of Trial 1. Trial 2 consisted of a 49-day feeding trial conducted under a similar initial stocking density and mean fish weight of 23.1 ± 0.8 g, as well as similar culture conditions, and tank systems as in Trial 1, with three replicates per dietary treatment in a randomized block design. Trial 3 consisted of a 56-day feeding trial performed in 500-L blue tanks under pilot-scale settings. Fish were initially stocked at a density of 25 individuals per tank with a mean body weight of 34.0 ± 1.2 g, with three replicates per dietary treatment in a randomized block design. Culture conditions were maintained as consistently as possible with those of the previous two feeding trials.

In all feeding trials, fish were fed experimental diets twice daily to apparent satiation throughout the study period, and mortality was recorded daily. Uneaten feed was monitored when applicable for feed intake and feed conversion ratio (FCR) calculations. All experimental procedures involving animals were performed in accordance with relevant institutional guidelines and regulations. This study was approved by the Institutional Animal Care and Use Committee (IACUC) under protocol number 202003-155A.

### Diet preparation

Experimental diets were formulated to evaluate microbial community-based single-cell protein (SCP) as a substitute for fishmeal in juvenile Asian seabass feeds. A fishmeal-based control diet (SCP0) and SCP-containing experimental diets were formulated by partially or completely replacing fishmeal with microbial community-based SCP while maintaining comparable crude protein and fat levels (Table 1). The detailed nutritional composition of the basal diets, including proximate composition, amino acid profiles, fatty acid composition, vitamin composition, and mineral composition, was previously characterized for Trial 1 (Santillan et al., 2024). The same basal formulation was maintained in Trials 2 and 3 to facilitate direct comparisons of experiments. In Trial 1, two dietary treatments were evaluated: SCP0 and SCP50, corresponding to 0% and 50% fishmeal replacement, respectively. Trial 2 evaluated SCP0, SCP50, and SCP100 diets, whereas Trial 3 evaluated SCP0, SCP10, SCP30, and SCP50 diets. Diet preparation followed the procedures previously described (Santillan et al., 2024). Briefly, dry feed ingredients were mixed thoroughly prior to the addition of liquid ingredients and water (20% v/v) using a stand mixer (Model 5 QT, KitchenAid, St Joseph, MI, USA). The resulting mash was passed through a meat grinder fitted with a 3-mm diameter die to produce moist feed strands, which were then manually broken into smaller pieces prior to drying overnight in a ventilated dryer at 45 °C. The dried feeds were subsequently stored in air-tight plastic containers at 4 °C until use.

**Table 1.** Composition of the fishmeal control (SCP0) and microbial community-based single-cell protein experimental diet replacing 50% of fishmeal protein (SCP50) used in the Asian seabass feeding trials^A^.

| Ingredient (% w/w) | SCP0 | SCP50 |
| --- | --- | --- |
| Fishmeal | 30.0 | 15.0 |
| SCP | 0.0 | 15.0 |
| Poultry by-product meal | 13.0 | 16.0 |
| Squid liver meal | 10.0 | 10.0 |
| Fish hydrolysate | 5.0 | 5.0 |
| Blood meal | 4.0 | 4.0 |
| Dried yeast | 3.0 | 3.0 |
| Fish oil | 8.0 | 8.0 |
| Wheat flour | 18.4 | 15.4 |
| Wheat gluten | 5.0 | 5.0 |
| Soy lecithin | 2.5 | 2.5 |
| Taurine | 0.5 | 0.5 |
| Ascorbic 2-phosphate 35% | 0.2 | 0.2 |
| Vitamin premix | 0.2 | 0.2 |
| Mineral premix | 0.2 | 0.2 |

<sup>A</sup> Diet formulations were maintained across trials to facilitate direct comparability among experiments. Adapted from (Santillan et al., 2024) under a CC BY 4.0 license.

### Growth performance and survival assessments

Fish were weighed at the beginning and end of each feeding trial to determine growth performance parameters. Feed intake was recorded at the tank level when applicable. Growth performance was evaluated by final body weight, mean weight gain (mean final weight – mean initial weight), percent weight gain (mean weight gain * mean initial weight^-1^ *100), specific growth rate (SGR = (ln_mean final weight_ – ln_mean initial weight_) * trial duration^-1^ *100), feed intake, FCR = mean feed intake * mean weight gain^-1^, and survival rate.

### Seabass tissue composition analysis

Whole-body proximate composition and essential amino acid analyses were performed on representative Asian seabass samples from the SCP0 and SCP50 dietary treatments, as these diets consistently achieved the highest growth performance and survival across the feeding trials. One pooled sample per dietary treatment (n = 1) was analysed by Merieux NutriSciences-AQ (Singapore) Pte. Ltd. according to standard analytical methods (AOAC, 1995). Analyses included determination of protein, lipid, ash, and essential amino acid compositions.

### SCP production

Microbial community-based SCP was produced using aerobic fermenters fed with soybean-processing side streams from a local soybean processing company (Mr Bean, Singapore), following methods previously described (Santillan et al., 2025; Santillan et al., 2024). For Trial 1, SCP was produced at lab scale using four 4-L sequencing batch reactors operated under non-axenic conditions with enriched microbial communities adapted to the substrate and operating conditions. Fermenters were operated on 12-h cycles with intermittent aeration, controlled temperature, and low DO concentrations to support biomass enrichment. For Trials 2 and 3, SCP was produced at pilot scale in a 100-L aerobic sequencing batch reactor operated for over 500 days. The pilot-scale system was inoculated with biomass derived from the lab-scale enrichments and fed soybean soaking side streams stored at low temperature prior to use. Due to the low nitrogen content of the side streams, nitrogen was supplemented externally as needed to maintain microbial growth. The fermenter was operated under low DO conditions and with hydraulic retention times ranging from 1 to 3 days. Biomass concentration was stabilized during later operational stages using membrane-based biomass retention approaches. Following cultivation, SCP biomass was harvested through sedimentation or membrane concentration, collected periodically, spray-dried, milled, and incorporated into the experimental aquafeeds.

### Nutritional characterization of feed ingredients

The nutritional composition of the microbial community-based SCP and commercial fishmeal used in the feeding trials was further characterized through amino acid and vitamin analyses. Protein content was quantified based on total amino acid composition using high-performance liquid chromatography (HPLC) following the procedure of Vethathirri et al. (2023). Essential amino acid profiles were normalized to total protein content to facilitate direct comparison between SCP and fishmeal ingredients. B-vitamin composition of SCP samples was determined by HPLC according to the method of Albawarshi et al. (2022).

### Microbial characterization of microbial community-based SCP

Genome-resolved metagenomic analyses were conducted on representative SCP batches associated with Asian seabass Trials 1 and 2 to characterize the microbial communities present within the SCP ingredient. To maximize genome recovery and assembly quality, representative samples were subjected to deep metagenomic sequencing using Oxford Nanopore long-read sequencing combined with Illumina short-read correction. Six samples (0.5 g each) were randomly collected from two 2-L bottles containing 300–500 g of microbial community-based SCP biomass, corresponding to triplicate samples from each bottle (n = 3 biological replicates per trial; 6 samples in total). DNA extraction was performed as previously described (Santillan et al., 2019).

Short-read metagenomic sequencing was performed using the Illumina HiSeq X platform in paired-end mode (2 × 150 bp), whereas long-read sequencing was conducted using the ONT PromethION platform. Illumina reads were quality-filtered and adapter-trimmed using BBDuk v39.01 (Bushnell, 2014), while ONT reads were filtered during basecalling using guppy v6.4.2. Hybrid metagenomic assemblies were generated using SPAdes v3.15.5 (Bankevich et al., 2012). Cleaned reads were subsequently mapped back to assembled contigs using Minimap2 v2.24 (Li, 2018), and alignment files were processed using SAMtools v1.17 (Li et al., 2009).

Metagenome-assembled genomes (MAGs) were recovered using complementary binning approaches including MetaBAT2 v2.15 (Kang et al., 2019), MaxBin2 v2.2.7 (Wu, Simmons, and Singer, 2016), and SemiBin2 v1.5 (Pan, Zhao, and Coelho, 2023), followed by refinement using MetaWRAP v1.4 (Uritskiy, DiRuggiero, and Taylor, 2018). Taxonomic classification of refined MAGs was performed using GTDB-Tk v1.7 (Chaumeil et al., 2020; Parks et al., 2022) against GTDB release 214. Genome quality assessment was conducted using CheckM v1.2.2 (Parks et al., 2015), and MAG quality categories were assigned according to the Minimum Information about Metagenome-Assembled Genomes (MIMAG) framework (Bowers et al., 2017). MAGs with completeness ≥ 50% and contamination ≤ 10% were retained for downstream analyses. Relative abundance estimation of recovered MAGs was performed using CoverM v0.6.1 (Aroney et al., 2025).

### Statistical analysis

Statistical analyses and data visualisation were performed in R version 4.4.1 using the packages *ggplot2* (v3.5.2), *rstatix* (v0.7.2), and *ggpubr* (v0.6.1). Growth performance and nutritional parameters were analysed separately within each trial. For comparisons involving two dietary treatments, Welch’s t-tests were used, whereas Welch’s analysis of variance (ANOVA) was applied when more than two dietary treatments were evaluated. To account for multiple testing across performance parameters, Welch’s ANOVA P-values were adjusted using the Benjamini-Hochberg false discovery rate (FDR) correction. When adjusted Welch’s ANOVA results remained significant (P_adj_ < 0.05), post hoc pairwise comparisons were performed using Games-Howell tests. For visualisation of performance trends across dietary inclusion levels SGR and FCR data from diets containing 0-50% SCP inclusion were plotted using locally estimated scatterplot smoothing (*loess*) regressions with 95% confidence intervals. Individual points represented replicate tanks colored according to feeding trial.

## Results and Discussion

### Microbial community-based SCP supports partial fishmeal replacement in juvenile Asian seabass diets

SCP maintained high survival and acceptable growth performance in juvenile Asian seabass at inclusion levels up to 50% fishmeal replacement in three independent feeding trials (Table 2; Figure 1). All treatments within the 0–100% SCP inclusion range achieved 100% survival. In Trials 1 and 3, no significant differences were detected in SGR, weight gain, or feed intake, whereas FCR differed significantly among treatments in Trial 3, with moderately higher values observed at higher inclusion levels. Regressions across the 0–50% SCP inclusion range similarly indicated stable SGR and modest increases in FCR with increasing SCP inclusion (Figure 1).

**Fig. 1.**
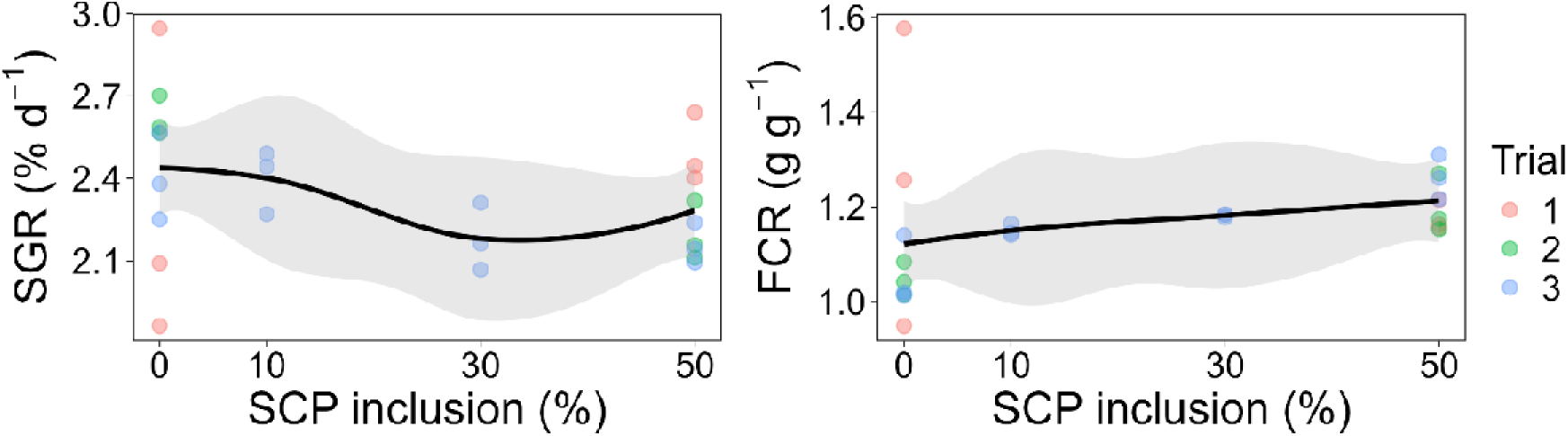
Specific growth rate (SGR) and feed conversion ratio (FCR) of juvenile *Lates calcarifer* as a function of dietary fishmeal replacement with microbial community-based single-cell protein (SCP). Data are from three independent feeding trials with 0 to 50% SCP inclusion. 0% inclusion corresponds to the control diet. Points represent individual tanks, colored according to trial. Black lines represent *loess* regressions fitted across all trials, and shaded areas indicate 95% confidence intervals.

**Table 2.**
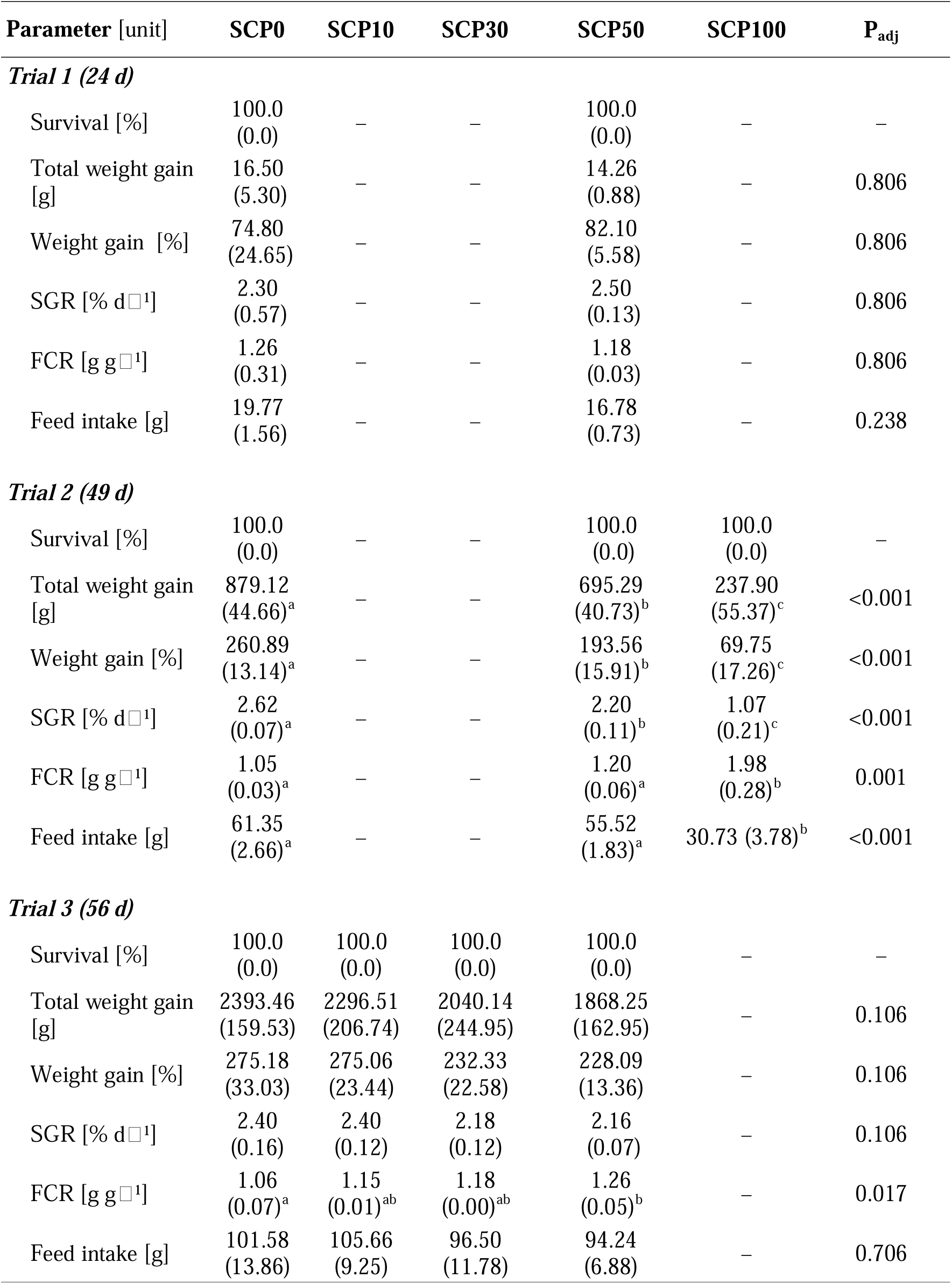
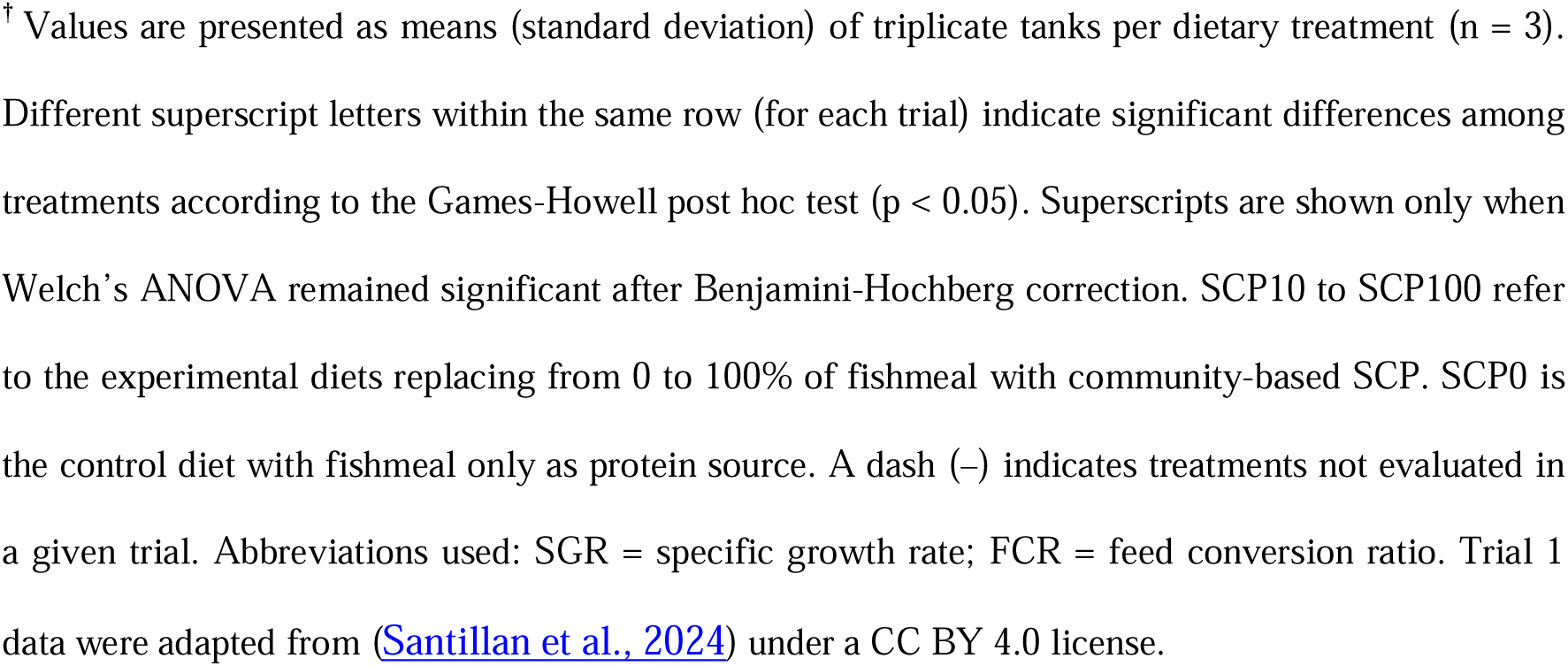
Growth performance of juvenile Asian seabass fed microbial community-based SCP diets in three independent feeding trials**^†^**.

| Parameter [unit] | SCP0 | SCP10 | SCP30 | SCP50 | SCP100 | P<sub>adj</sub> |
| --- | --- | --- | --- | --- | --- | --- |
| **Trial 1 (24 d)** |  |  |  |  |  |  |
| Survival [%] | 100.0<br>(0.0) | — | — | 100.0<br>(0.0) | — | — |
| Total weight gain [g] | 16.50<br>(5.30) | — | — | 14.26<br>(0.88) | — | 0.806 |
| Weight gain [%] | 74.80<br>(24.65) | — | — | 82.10<br>(5.58) | — | 0.806 |
| SGR [% d<sup>-1</sup>] | 2.30<br>(0.57) | — | — | 2.50<br>(0.13) | — | 0.806 |
| FCR [g g<sup>-1</sup>] | 1.26<br>(0.31) | — | — | 1.18<br>(0.03) | — | 0.806 |
| Feed intake [g] | 19.77<br>(1.56) | — | — | 16.78<br>(0.73) | — | 0.238 |
| **Trial 2 (49 d)** |  |  |  |  |  |  |
| Survival [%] | 100.0<br>(0.0) | — | — | 100.0<br>(0.0) | 100.0<br>(0.0) | — |
| Total weight gain [g] | 879.12<br>(44.66)<sup>a</sup> | — | — | 695.29<br>(40.73)<sup>b</sup> | 237.90<br>(55.37)<sup>c</sup> | <0.001 |
| Weight gain [%] | 260.89<br>(13.14)<sup>a</sup> | — | — | 193.56<br>(15.91)<sup>b</sup> | 69.75<br>(17.26)<sup>c</sup> | <0.001 |
| SGR [% d<sup>-1</sup>] | 2.62<br>(0.07)<sup>a</sup> | — | — | 2.20<br>(0.11)<sup>b</sup> | 1.07<br>(0.21)<sup>c</sup> | <0.001 |
| FCR [g g<sup>-1</sup>] | 1.05<br>(0.03)<sup>a</sup> | — | — | 1.20<br>(0.06)<sup>a</sup> | 1.98<br>(0.28)<sup>b</sup> | 0.001 |
| Feed intake [g] | 61.35<br>(2.66)<sup>a</sup> | — | — | 55.52<br>(1.83)<sup>a</sup> | 30.73 (3.78)<sup>b</sup> | <0.001 |
| **Trial 3 (56 d)** |  |  |  |  |  |  |
| Survival [%] | 100.0<br>(0.0) | 100.0<br>(0.0) | 100.0<br>(0.0) | 100.0<br>(0.0) | — | — |
| Total weight gain [g] | 2393.46<br>(159.53) | 2296.51<br>(206.74) | 2040.14<br>(244.95) | 1868.25<br>(162.95) | — | 0.106 |
| Weight gain [%] | 275.18<br>(33.03) | 275.06<br>(23.44) | 232.33<br>(22.58) | 228.09<br>(13.36) | — | 0.106 |
| SGR [% d<sup>-1</sup>] | 2.40<br>(0.16) | 2.40<br>(0.12) | 2.18<br>(0.12) | 2.16<br>(0.07) | — | 0.106 |
| FCR [g g<sup>-1</sup>] | 1.06<br>(0.07)<sup>a</sup> | 1.15<br>(0.01)<sup>ab</sup> | 1.18<br>(0.00)<sup>ab</sup> | 1.26<br>(0.05)<sup>b</sup> | — | 0.017 |
| Feed intake [g] | 101.58<br>(13.86) | 105.66<br>(9.25) | 96.50<br>(11.78) | 94.24<br>(6.88) | — | 0.706 |

<sup>†</sup> Values are presented as means (standard deviation) of triplicate tanks per dietary treatment (n = 3). Different superscript letters within the same row (for each trial) indicate significant differences among treatments according to the Games-Howell post hoc test ( $p < 0.05$ ). Superscripts are shown only when Welch's ANOVA remained significant after Benjamini-Hochberg correction. SCP10 to SCP100 refer to the experimental diets replacing from 0 to 100% of fishmeal with community-based SCP. SCP0 is the control diet with fishmeal only as protein source. A dash (–) indicates treatments not evaluated in a given trial. Abbreviations used: SGR = specific growth rate; FCR = feed conversion ratio. Trial 1 data were adapted from (Santillan et al., 2024) under a CC BY 4.0 license.

In Trial 2, complete fishmeal replacement (SCP100) resulted in significantly lower feed intake, weight gain, and SGR relative to SCP0 and SCP50 (Table 2). These results suggest that performance reductions at complete replacement were more likely associated with palatability or formulation constraints than broad limitations in nutrient utilization. Similar reductions at high microbial protein inclusion levels have been associated with altered sensory properties, amino acid imbalances, elevated nucleic acid content, or reduced attractability in aquaculture feeds (Li et al., 2024; Mugwanya, Kimera, and Sewilam, 2026). The experimental diets already included blood meal and fish hydrolysate, indicating that additional formulation optimization may still be required at very high inclusion levels.

The SCP biomass exhibited nutritionally relevant protein, amino acid, and B-vitamin compositions (Table 3). Across trials, SCP inclusion levels up to 50% fishmeal replacement maintained high survival and generally stable performance in juvenile Asian seabass, supporting a reproducible pattern of partial fishmeal replacement without major reductions in overall performance. Although SCP100 reduced growth-related metrics, survival remained high, indicating that complete fishmeal replacement did not compromise fish viability under the conditions tested.

**Table 3.**
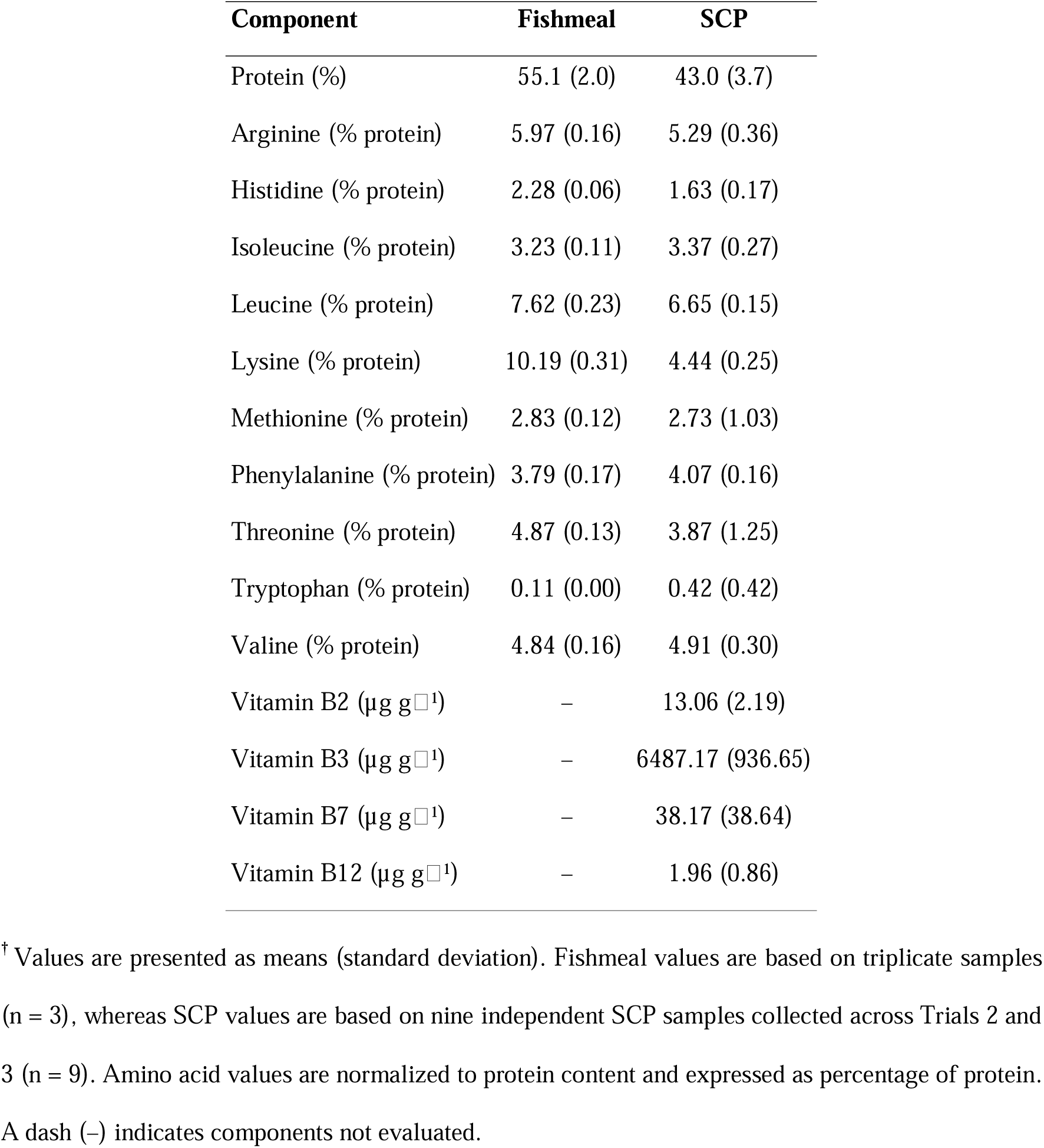
Total amino acid-derived protein content, essential amino acid composition, and B-vitamin composition of fishmeal and microbial community-based SCP used in Asian seabass feeding trials**^†^**.

| Component | Fishmeal | SCP |
| --- | --- | --- |
| Protein (%) | 55.1 (2.0) | 43.0 (3.7) |
| Arginine (% protein) | 5.97 (0.16) | 5.29 (0.36) |
| Histidine (% protein) | 2.28 (0.06) | 1.63 (0.17) |
| Isoleucine (% protein) | 3.23 (0.11) | 3.37 (0.27) |
| Leucine (% protein) | 7.62 (0.23) | 6.65 (0.15) |
| Lysine (% protein) | 10.19 (0.31) | 4.44 (0.25) |
| Methionine (% protein) | 2.83 (0.12) | 2.73 (1.03) |
| Phenylalanine (% protein) | 3.79 (0.17) | 4.07 (0.16) |
| Threonine (% protein) | 4.87 (0.13) | 3.87 (1.25) |
| Tryptophan (% protein) | 0.11 (0.00) | 0.42 (0.42) |
| Valine (% protein) | 4.84 (0.16) | 4.91 (0.30) |
| Vitamin B2 (µg g<sup>-1</sup>) | – | 13.06 (2.19) |
| Vitamin B3 (µg g<sup>-1</sup>) | – | 6487.17 (936.65) |
| Vitamin B7 (µg g<sup>-1</sup>) | – | 38.17 (38.64) |
| Vitamin B12 (µg g<sup>-1</sup>) | – | 1.96 (0.86) |

<sup>†</sup> Values are presented as means (standard deviation). Fishmeal values are based on triplicate samples (n = 3), whereas SCP values are based on nine independent SCP samples collected across Trials 2 and 3 (n = 9). Amino acid values are normalized to protein content and expressed as percentage of protein. A dash (–) indicates components not evaluated.

### SCP inclusion outcomes in Asian seabass diets depend strongly on production system and formulation

The study demonstrated consistently high survival and generally stable performance at up to 50% fishmeal replacement in three independent trials using juvenile fish and two production scales (120- and 500-L fish tanks). Complete fishmeal replacement did not affect fish viability but reduced feed intake and growth, indicating that limitations remain at very high inclusion levels. Collectively, these findings indicate that SCP should not be treated as a single ingredient category, as microbial composition, production process, fish developmental stage, biomass processing, and dietary formulation jointly influence nutritional and physiological outcomes.

Other Asian seabass studies have demonstrated highly variable outcomes depending on SCP source, production system, dietary formulation, and fish developmental stage (Mugwanya, Kimera, and Sewilam, 2026). Methanotrophic bacterial SCP replacing 25–50% of fishmeal in barramundi fry induced poor growth, reduced survival, enteropathy, liver pathology, and gut microbiome dysbiosis associated with increased *Aeromonas* abundance (Samsing et al., 2024). In contrast, other methanotrophic bacterial ingredient studies reported more favorable outcomes on growth performance, feed utilization, intestinal physiology, and gut microbial composition in barramundi (Woolley et al., 2023). Enriched purple phototrophic bacterial biomass also supported partial fishmeal replacement, although complete replacement reduced growth and increased FCR (Delamare-Deboutteville et al., 2019). Brewery-derived bacterial protein meals additionally demonstrated favorable digestibility coefficients and digestive enzyme responses in barramundi juveniles (Le Boucher et al., 2025). Collectively, these studies further support that physiological responses to SCP depend strongly on ingredient origin, microbial composition, processing, dietary formulation, and fish developmental stage. Our study additionally combined multiple feeding trials conducted across different production scales with nutritional characterization and genome-resolved metagenomic analyses of the SCP ingredient.

### Partial fishmeal replacement maintained whole-body nutritional composition

Whole-body proximate composition and essential amino acid profiles of fish fed SCP0 and SCP50 diets were broadly similar (Table 4). Fish receiving SCP50 exhibited protein, lipid, and ash contents comparable to those of fish fed the fishmeal-based control diet, while amino acid profiles including lysine, methionine, leucine, valine, and threonine remained generally consistent between treatments. Maintenance of whole-body composition under partial fishmeal replacement is relevant because substantial nutritional imbalances may affect both fish performance and product quality. The relatively stable amino acid composition observed here is consistent with previous reports describing nutritionally relevant amino acid profiles in food-processing side stream-derived SCP systems (Vethathirri et al., 2023; Vethathirri et al., 2025; Xu et al., 2026). Although body composition analyses were based on pooled samples, these findings further support the suitability of SCP as a partial fishmeal substitute in juvenile Asian seabass diets.

**Table 4.** Whole-body nutritional composition and essential amino acid profiles of juvenile Asian seabass (freeze-dried basis) fed the fishmeal control (SCP0) or microbial community-based SCP diet (SCP50)**^†^**.

| Component | SCP0 (%) | SCP50 (%) | Method<sup>‡</sup> |
| --- | --- | --- | --- |
| *Proximate composition* |  |  |  |
| Protein | 54.3 | 55.8 | A |
| Lipid | 34.8 | 31.6 | B |
| Ash | 14.4 | 12.9 | C |
| *Essential amino acid composition* |  |  |  |
| Histidine | 0.63 | 0.65 | D |
| Threonine | 1.00 | 1.02 | D |
| Valine | 1.81 | 1.88 | D |
| Methionine | 1.22 | 1.25 | D |
| Tryptophan | 0.34 | 0.39 | E |
| Phenylalanine | 1.48 | 1.52 | D |
| Isoleucine | 1.23 | 1.26 | D |
| Leucine | 2.60 | 2.70 | D |
| Lysine | 3.09 | 3.19 | D |

<sup>†</sup> Values are based on a single pooled feed sample per dietary treatment (n = 1).
<sup>‡</sup> A, AsureQuality Method 6503s, Kjeldahl block digestion; B, AsureQuality Method 4988, Gravimetric-SBR; C, AsureQuality Method 4151, Gravimetric; D, Modified from Analytical Biochemistry 178 (1989); E, AsureQuality Method 5707, HPLC.

### Genome-resolved metagenomics revealed diverse microbial taxa associated with SCP production

Metagenomic assembly and binning recovered numerous medium- and high-quality MAGs representing diverse bacterial taxa associated with the SCP production system (Table 5, Table S1). Recovered taxa included members affiliated with *Acidipropionibacterium*, *Lactococcus, Limosilactobacillus, Bifidobacterium, Paracoccus,* and members of the family *Rhodobacteraceae*, many of which have previously been associated with fermentation processes, vitamin biosynthesis, organic acid production, or metabolite production. Previous characterization of side stream-derived SCP similarly identified taxa with potential probiotic or vitamin-producing capabilities, including *Lactococcus, Weissella*, and *Acidipropionibacterium* (Vethathirri et al., 2023). Compared with marker gene-based community profiling approaches, genome-resolved metagenomics additionally provides genome-level resolution that can improve characterization of microbial community composition and support future investigations into metabolic and functional potential (Mirete et al., 2025; Neshat et al., 2024). The recovery of diverse MAGs across independent SCP production systems further supports the feasibility of generating nutritionally consistent microbial community-based biomass using non-axenic cultivation approaches. Compared with axenic systems, microbial community-based production systems may offer advantages in substrate flexibility, operational robustness, and scalability, properties that have been associated with complex microbial communities in environmental biotechnology systems (Santillan, Neshat, and Wuertz, 2025; Vethathirri, Santillan, and Wuertz, 2021).

**Table 5.** Quality assessment of metagenome-assembled genomes (MAGs) recovered from metagenomes associated with Asian seabass Trials 1 and 2 (n = 3 biological replicates per trial; 6 samples in total).

| Quality<sup>†</sup> | Completeness<sup>‡</sup> | Contamination | Count |
| --- | --- | --- | --- |
| Putative high-quality | 90 – 100 % | <5 % | 243 |
| Medium-quality | 50 – 90 % | <10% | 304 |
| Total |  |  | 547 |

<sup>†</sup> Quality assigned based on the Minimum Information about Metagenome-Assembled Genomes (MIMAG) framework (Bowers et al., 2017).
<sup>‡</sup> CheckM completeness estimates the percentage of expected lineage-specific single-copy marker genes recovered in a MAG, indicating how complete the genome is (Parks et al., 2015).

Microbial community-based SCP systems may additionally provide functional properties extending beyond crude protein replacement through the retention of microbial metabolites and structural cellular components with potential postbiotic activity. Previous studies reported enhanced stress tolerance, immune responses, or survival in shrimp fed purple non-sulfur bacterial biomass or microbial community-based SCP systems (Alloul et al., 2021; Santillan et al., 2025). Although the present study focused primarily on nutritional performance and microbial community characterization, the recovered MAG dataset provides a foundation for future genome-resolved investigations into metabolic pathways and potential functional properties associated with microbial community-based SCP systems, including possible interactions with host physiology and gut microbiomes.

### Translational relevance and study limitations

The present study demonstrated reproducible performance of SCP across independent feeding trials and production scales despite the variability commonly reported in microbial protein studies (Mugwanya, Kimera, and Sewilam, 2026). The use of microbial community-based production systems additionally supports the feasibility of utilizing nutrient-rich industrial side streams without highly controlled sterile production processes, potentially improving production sustainability and cost efficiency while reinforcing the translational potential of side stream-derived SCP within circular bioeconomy strategies for aquaculture feed production.

Several limitations should nevertheless be acknowledged. Although the present trials were comparable to or longer than most published Asian seabass SCP feeding studies, they did not cover a full grow-out cycle to harvest size. Whole-body composition analyses were conducted using pooled samples, and host gut microbiome responses, histology, immune biomarkers, and pathogen challenge outcomes were not evaluated directly. The study also focused primarily on juvenile fish performance under controlled experimental conditions rather than commercial grow-out systems. Additional long-term feeding and genome-resolved functional studies would help further clarify the nutritional and functional potential of microbial community-based SCP systems in aquaculture feeds.

## Conclusions

This study demonstrated that microbial community-based SCP could reproducibly replace up to 50% of fishmeal in juvenile Asian seabass (*Lates calcarifer*) diets in three independent feeding trials involving different growth periods and tank volumes. Partial fishmeal replacement maintained high survival, generally stable growth performance, and comparable whole-body composition relative to conventional fishmeal-based diets. Although complete fishmeal replacement reduced feed intake and growth, the performance limitations at very high SCP inclusion levels were likely associated primarily with feed formulation and palatability constraints rather than impaired nutrient utilization. Genome-resolved metagenomic analyses further revealed diverse microbial taxa associated with the SCP ingredient, supporting the feasibility of microbial community-based production systems for aquaculture feed applications. Collectively, these findings demonstrate reproducible partial fishmeal replacement across independent trials and production scales while supporting the feasibility of microbial community-based SCP systems for sustainable finfish aquaculture. The integration of nutritional characterization, production-scale validation, and genome-resolved microbial analyses further provides a framework for evaluating microbial community-based SCP systems as multifunctional aquafeed ingredients.

## Data availability

MAG quality assessment data supporting this study are provided in Table S1.

## Author Contributions

ES, PLL, FY, DCPS, and SW conceived the studies and designed the experiments. SW and DCPS secured funding for the study. XH, RSV, and ES produced the microbial protein. FY conducted Trial 1, while PLL conducted Trials 2 and 3 and associated wet-lab work, including feed preparation, animal husbandry, sample collection, and data reporting. PLL compiled and organized the data from Trials 2–3. ES coordinated the SCP molecular work and chemical analyses, performed the statistical analyses and data visualisation, interpreted the results, and integrated datasets across trials. SAN designed the metagenomic sequencing strategy and conducted the bioinformatics analyses. ES wrote the first draft of the manuscript. All authors reviewed and approved the final manuscript.

